# Sensitivity in binding free energies due to protein reorganization

**DOI:** 10.1101/066621

**Authors:** Nathan M. Lim, Lingle Wang, Robert Abel, David L. Mobley

**Affiliations:** Department of Pharmaceutical Sciences, University of California—Irvine, Irvine, California 92697, United States; Schrödinger, Inc., 120 West 45th Street, New York, New York 10036, United States

## Abstract

Tremendous recent improvements in computer hardware, coupled with advances in sampling techniques and force fields, are now allowing protein-ligand binding free energy calculations to be routinely used to aid pharmaceutical drug discovery projects. However, despite these recent innovations, there are still needs for further improvement in sampling algorithms to more adequately sample protein motion relevant to protein-ligand binding. Here, we report our work identifying and studying such clear and remaining needs in the apolar cavity of T4 Lysozyme L99A. In this study, we model recent experimental results that show the progressive opening of the binding pocket in response to a series of homologous ligands.^1^ Even while using enhanced sampling techniques, we demonstrate that the predicted relative binding free energies (RBFE) are sensitive to the initial protein conformational state. Particularly, we highlight the importance of sufficient sampling of protein conformational changes and demonstrate how inclusion of three key protein residues in the ‘hot’ region of the FEP/REST simulation improves the sampling and resolves this sensitivity.

## 1 Introduction

Proteins play a central role in many biological processes by modulating many key signaling pathways. It is unsurprising to find that proteins make up the vast majority of pharmaceutical drug targets. Many small-molecules on the market today induce their therapeutic effect by modulating the proteins’ biological activity through binding.^2–4^ Thus, optimization of protein-ligand binding affinities is a central goal in early pharmaceutical drug design projects.

Recent advancements in technology and computational chemistry have led to increasing use of computer-aided drug design techniques to assist in lead optimization in pharmaceutical drug discovery. Since accuracy and reliability of the approach being used is critical for success, it is here where the most rigorous of methods like free energy calculations can be applied in the prediction of protein-ligand binding affinities. Using alchemical methods like free energy perturbation (FEP), thermodynamic integration, and *λ*-dynamics on molecular dynamics (MD) simulations, the difference in binding free energies between two ligands can be computed in a robust and accurate manner. By computing relative binding free energies, much of the computational cost and difficulties of absolute binding free energy calculations is avoided.^5–11^ Other advancements in forcefields, sampling algorithms, and emergence of GPU computing have considerably improved the accuracy and robustness of alchemical calculations. The recent development of FEP+, a fully automated alchemical protocol implemented with modern methodologies, reduces overall workload and the potential for human error in setup and analysis of these types of calculations. Using this protocol, previous studies^12^ have demonstrated it to yield highly accurate free energy predictions in a wide range of pharmaceutically relevant protein targets and ligands. The accuracy and reliability of FEP+ makes it a very powerful tool in the hands of medicinal chemists for efficient optimization of lead compounds.

Although the previous report on FEP+^12^ demonstrates its robustness, the primary concern focuses on the size of the ligand perturbation or the initial ligand pose and how these factors impact the accuracy of calculated free energies.^13,14^ Careful consideration of effects arising from protein conformational changes have generally received less attention due to the difficulty in addressing the sampling challenges encountered when simulating proteins. The difficulty in protein sampling lies primiarly in the timescales required to adequately capture a complete biomolecular event. Timescales of such events can range from nanoseconds to milliseconds (or even seconds).^15^ In MD simulations, the system evolves in time through series of short time steps (i.e. 2 fs) by repeatedely computing the forces on each individual atom in accordance to Newton’s laws of motion at each time step. Not only must a computer perform numerous calculations at each time step, but it must repeat this process an enour-mous amount of times to generate a trajectory that is on timescale for your biological process of interest.^16^ Furthering the difficulty, with using ‘vanilla’ MD simulations, the system will more than likely remain kinetically trapped in an energy minima, thereby preventing any further exploration of the protein conformational space and obtaining proper sampling. Some computational studies have shown even small simple changes in the protein—like a side-chain rotamer flipping—can cause large errors in affinity predictions if not properly sampled.^17–20^ Overcoming these high energy barriers is no easy feat; whereby a number of distinct approaches like metadynamics,^21^ accelerated MD,^22^ and temperature accelerated MD^23^ aim to solve this problem of kinetic traps but may fall short in some cases.^24^ Thus, careful sampling of protein conformational changes is critical for accurate and reliable affinity predictions.

In the previous FEP+ study,^12^ other than simply restricting ourselves to congeneric series with multiple reported crystal structures, no effort was made to either include or exclude cases where the protein or ligand binding mode may reorganize. However, it is entirely possible some of the outliers reported in that work may be due to protein conformational reorganization. The testing of the FEP+ technology on cases where protein reorganization has been well-characterized experimentally may provide an opportunity to probe potential pathological cases, and further provides an opportunity to discover how to further improve the technology. While some studies suggest protein reorganization on ligand binding can be relatively common (i.e. as reviewed by^25^), it is far from well understood how often this can be a challenge for binding prediction. Regardless, it is clear that protein-ligand binding can be highly complex, and multiple stable protein conformations may be populated even for the same ligand.^1,26,27^ Thus, there is a great need to understand how protein conformational changes and the choice of starting protein conformation may affect the reliability of free energy predictions.

Here, we apply the now-standard FEP protocol used in,^12^ which utilizes replica exchange with solute tempering (FEP/REST),^28^ to a very simple model binding site in an engineered mutant of T4 lysozyme (L99A). In this mutant, the L99A mutation creates a small apolar binding site that has been studied extensively experimentally^29–32^ and computationally by docking^1,33–35^ and free energy methods.^18,28,36–41^ Recent studies on this binding site have found that the protein adopts three discrete conformations in response to ligand binding.^1^ Through a series of eight congeneric ligands (Fig 1f), each growing by addition of a single methyl, the protein responds by a single helix rearrangement to accommodate the growing ligand (Fig 1). As the ligand grows, the binding cavity was observed to incrementally reorganize into three discrete conformations which we will refer to as the closed (Fig 1c), intermediate(Fig 1d), and open states (Fig 1e). Using the FEP/REST protocol and the aforementioned homologous ligand series, we calculate relative protein-ligand binding affinities between ligands that occupy different discrete protein conformational states. In this study—while using the implemented default FEP/REST protocol—we demonstrate how the kinetically distinct protein states and structural rearrangement affects the accuracy and reliability of our predicted relative binding affinities. Further, we illustrate the importance of sufficient sampling of protein conformational changes and how modification of the REST region can potentially address this issue.

**Figure 1:**
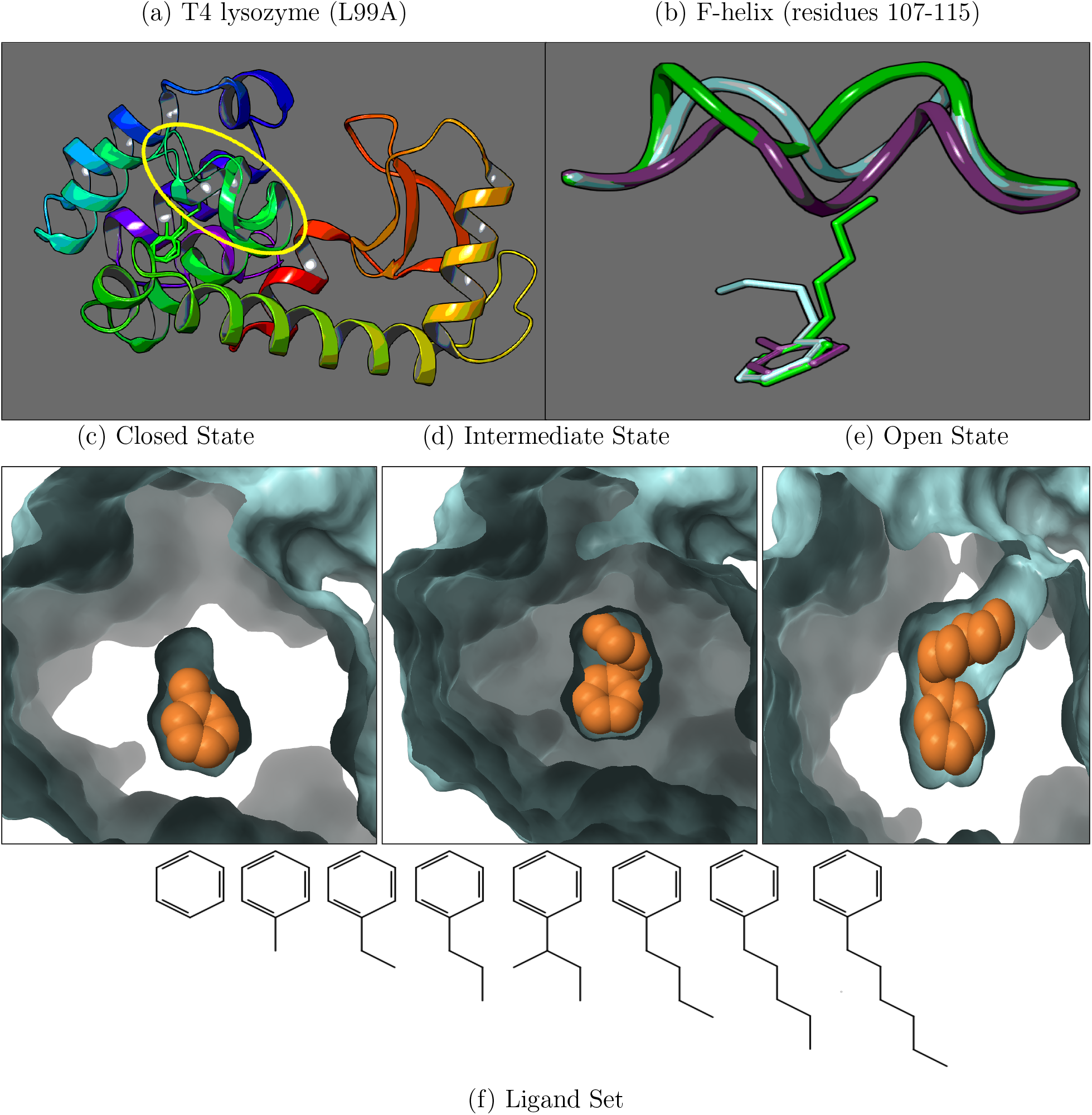
In Fig.1a are overlaid cartoon representations of T4 lysozyme using the protein-ligand bound crystal structures of the toluene(4W53), butylbenzene(4W57), and hexylbenzene(4W59) complexes where the F-helix region is highlighted in yellow. Fig.1b illustrates a closer view of strictly the F-helix in the closed(purple), intermediate(cyan), and open(green) conformational states and their corresponding ligands as found in the crystal structure. Below are the molecular surface representations of the binding cavity in the closed(1c), intermediate(1d), and open(1e) states with the ligands represented in an orange space-filling model. In Fig.1f is the full series of congeneric ligands used in this study; their protein state occupancies can be found in Table 1. Images were created using Maestro.^42^

## 2 Methods

### Protein/Ligand preparation

All proteins were prepared and aligned in Maestro^42^ using the ‘Protein Preparation Wizard’^43–47^ tool and with the following settings enabled (as they appear in the Maestro GUI menu):

- Preprocess: Assign bond orders, Add hydrogens, Create zero-order bonds to metals, Create disulfide bonds, Cap termini, Delete waters beyond 5Å from het groups
- Refine: Sample water orientations, Use PROPKA pH: 7.0, Remove waters with less than 3 H-bonds to non-waters, and restrained minimization.

Protein structures were taken from PDBs: 4W52, 4W53, 4W54, 4W55, 4W56, 4W57, 4W58, and 4W59 corresponding to protein-ligand bound structures of benzene, toluene, ethylbenzene, propylbenzene, sec-butylbenzene, butylbenzene, pentylbenzene, and hexylbenzene, respectively.^1^ Each simulation starts from either the protein closed state (PDB:4W52) or the open state (PDF:4W59). Using LigandFEP, our system preparation follows a similar workflow to the tutorial.^48^ Generally, two options were taken:

(1a) If the simulation starts from the protein closed state, the benzene crystal position was used as a reference for fragment building (PDB:4W52).
(1b) The corresponding ligand in the transformation was built by duplicating benzene in place and adding methyl groups.
(2a) If the simulation starts from the protein open state, the hexylbenzene crystal position was used as reference for fragment building (PDF:4W59).
(2b) The corresponding ligand in the transformation was built by duplicating hexylbenzene in place and deleting methyl groups.

Ligand tail fragments were added using the Build/Fragments toolbar in Maestro and were not overlaid or docked. As the ligand tails were built, bonds were manually rotated so that the tail was oriented in a similar manner as in their corresponding crystal structure. Following, the newly added atoms in the tail were locally minimized while leaving the core in its initial position. This was done in an attempt to correct bond angles and minimize the core RMSD, which LigandFEP uses to determine the core atoms between the two ligands.

### Classification of alchemical transformations and color coding

Here, we classified ligands based on the primary protein conformation (closed, intermediate, or open) the ligand occupies from the experimental studies (Table 1). To be explicit, the set of closed ligands refers to benzene, toluene, ethylbenzene, and propylbenzene; (sec-)butylbenzene for intermediate; and pentyl/hexylbezene for open ligands. The various protein states and ligands are then assigned a color accordingly: purple for closed, cyan for intermediate, and green for the open state.

**Table 1:** Experimental protein-ligand occupancies and affinities^*a*^

| PDB | Ligand | C | I | O | $\Delta G_{exp}$ | $\sigma_{exp}$ | Set |
| --- | --- | --- | --- | --- | --- | --- | --- |
| 4W52 | benzene | 90 | - | - | -5.19 | 0.16 | C |
| 4W53 | toluene | 80 | 20 | - | -5.52 | 0.04 | C |
| 4W54 | ethylbenzene | 50 | 50 | - | -5.76 | 0.07 | C |
| 4W55 | propylbenzene | 60 | 40 | - | -6.55 | 0.02 | C |
| 4W56 | sec-butylbenzene | 40 | 60 | - | N/A | - | I |
| 4W57 | butylbenzene | 10 | 60 | 30 | -6.70 | 0.02 | I |
| 4W58 | pentylbenzene | 30 | - | 70 | N/A | - | O |
| 4W59 | hexylbenzene | 30 | - | 70 | N/A | - | O |

<sup>a</sup> Listed are the PDB codes of the corresponding protein-ligand complex crystal structures and the ligand occupancies<sup>1</sup> for the closed(C), intermediate(I), and open(O) protein conformations by percentage. Under the ‘Set’ column, shaded in grey, is each ligand’s primary state of occupancy which we use to categorize our alchemical transformation sets. Binding affinities for pentylbenzene and hexylbenzene were inaccessible due to solubility limits.<sup>1</sup> Experimental binding free energies and their uncertainties are in units of kcal/mol.<sup>31</sup>

Our sets of alchemical transformations consisted of 3 groups: ‘closed-intermediate’, ‘closed-open’, and an ‘experimental’ ligand set. The first two sets were classified based on the expected conformational change that would result from the alchemical transformation. For example, the alchemical transformation of benzene to butylbenzene falls into closed-intermediate while benzene to hexylbenzene is a closed-open transformation. In our experimental set, we perform all possible combinations of transformations for ligands with available experimental binding affinities (Table 1). Ligands with available experimental binding affinities consisted of benzene, toluene, ethylbenzene, propylbenzene, and butylbenzene. This gives a total of 26 alchemical transformations in this study, 8 from ‘closed-intermediate’, 8 from ‘closed-open’, and 10 from the experimental set.

### FEP protocols

Using the Schrödinger application suite (release 2015-3),^49^ we utilize a similar protocol to FEP+^12^ called LigandFEP.^48^ FEP+ is a fully automated work flow that plans perturbation pathways based a variant of LOMAP^50^ mapping algorithm which uses the maximum common substructure (MCS) between any pair of compounds. LigandFEP is an academic toolkit that generates the configuration files to perform the free energy calculation but is limited in the sense that the user must plan each perturbation path instead. Both LigandFEP and FEP+ use the same Desmond relaxation protocol and the FEP/REST methodology.^28,41,51,52^ By utilizing LigandFEP, we demonstrate LigandFEP can be a powerful tool for academics as it strictly differs only in the level of automation.

### REST region selection

In this study, by default, only heavy atoms in the ligand were included in the REST region unless specified otherwise. Further details on the temperature profile and how the REST region is normally selected can be found in previous studies^12,28^ in the supporting information. Simulations that included protein heavy atoms in the REST region are referred to with the ‘pREST’ label, where selection of the particular residues is described as follows.

Based on visual inspection of our molecular dynamics simulations and considering the F-helix spans residues 107-115, we selected residues Glu108, Val111, and Gly113 to include into the REST region (Fig 2a). Glu108 sits near the start of the helix which appears as a hinge point for the opening and closing of the binding cavity (Fig 2b). Following, Val111 appears in the middle of the helix and was observed to undergo the largest motion during protein conformational changes (Fig 2c). Gly113 was included in order to collectively have hot regions approximately at the start, middle and end points of the helix.

**Figure 2:**
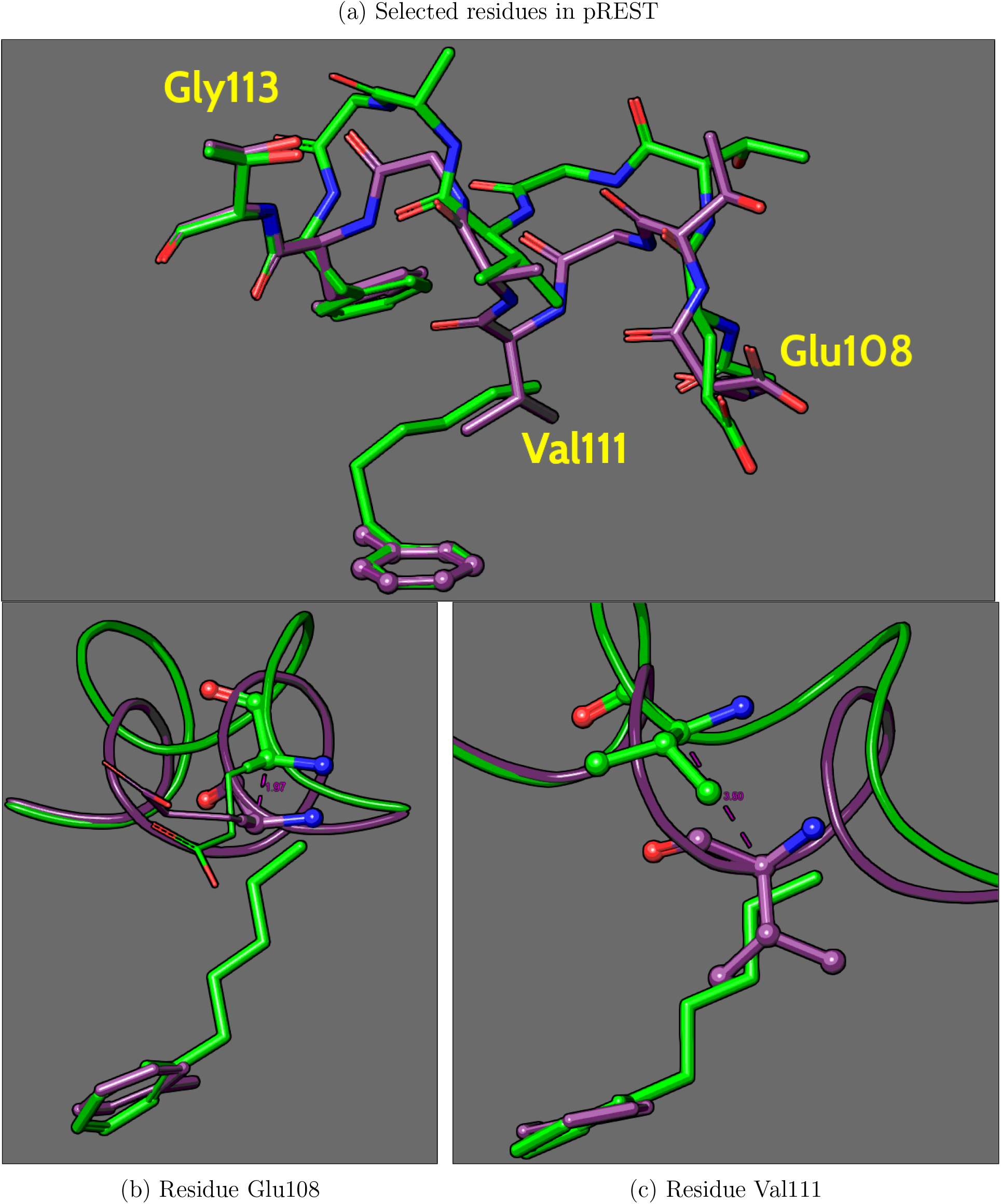
Fig. 2a highlights the 3 residues selected to be included into the REST region for ‘pREST’ simulations. Fig. 2b and Fig. 2c illustrates the motion of the *C_α_*’s during the transition between the protein closed and open states, where the *C_α_* in Glu108 undergoes a motion of approximately 2Å, while in Valll the *C_α_* move approximately 3.5Å.

### Simulation details

Desmond^53–56^ simulation protocols have been described previously in the supporting information^12^ or can be found in greater detail in the Desmond User Manual.^57^ The relaxation protocol begins with a simulation where solute molecules are restrained to their initial positions while minimizing using a Brownian dynamics NVT integrator for 100ps, followed by 12ps simulations at 10K with a NVT ensemble and then a NPT ensemble using the Langevin method.^58^ Next is a 24ps simulation followed by a final 240ps simulation with solute molecules unrestrained, both are carried at room temperature with a NPT ensemble using Langevin. Production simulations with the default REST region were ran for the default setting of 5ns. Initial ‘pREST’ production simulations were also simulated for 5ns and then were carried out to a length of up to 55ns for all closed-open transformations. For closed-intermediate transformations, we extended ‘pREST’ production simulations up to 25ns, only for cases that were far from convergence with the 5ns simulation time. Here, we use the final 15ns for closed-open simulations and the final 10ns for closed-intermediate to calculate our final free energies, discarding the initial time as additional equilibration time. FEP/REST simulations were run on four GeForce GTX Titan Black GPUs using the Desmond/GPU engine with the recently developed OPLS3^59^ forcefield parameters.

### Calculation of free energies and measurement of inconsistency

Throughout this study, we measure the inconsistency (ΔΔ*G_ε_n__*) between the final calculated free energies between simulations that start from the protein closed state (ΔΔ*G*_C_n__) versus the protein open state (ΔΔ*G*_O_n__) by simply taking the difference.

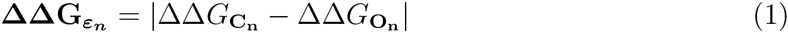

Then we compute the overall inconsistency—referred to as the ‘Root-Mean-Square-Inconsistency’ (RMSI)—for each set of alchemical transformations. The RMSI is calculated by using the differences (ΔΔ*G_ε_n__*) obtained from the comparisons between protein open and closed simulations.

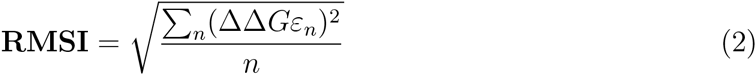

Similarly, we compute the ‘Root-Mean-Square-Error’(RMSE) when comparing with experimental free energies for both simulations starting from the protein closed and open state.

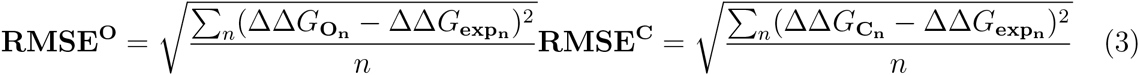

Calculated free energies were determined using the Bennett acceptance ratio^60^ (BAR) with error estimations using both bootstrapping and BAR analytical error prediction.^61^ Hysteresis around closed thermodynamic cycles and best estimates of the free energies with their errors were calculated using the cycle closure algorithm discussed in a previous publication.^28^

### Determining the protein conformation state using RMSD

In this study, we determine the state of the protein by computing the ‘Root-Mean-Square-Deviation’ (RMSD) of the protein backbone atoms spanning the F-helix relative to their positions found in the closed (PDB:4W53), intermediate (PDB:4W57), and open (PDB:4W59) crystal structures (Fig 1b). The set of RMSDs—that is, the RMSD relative to the closed, intermediate, and open states—is computed at each frame over the course of the entire simulation. Then, we use the protein conformational state with the lowest RMSD to correspondingly color each time point in our analyses of ‘RMSD/time’ and ‘Color maps’. Again, we use purple to denote the protein closed state, cyan for the intermediate state, and green for the open state. Here, we use VMD^62,63^ to align and compute the RMSD of our Desmond trajectories relative to crystal structures. Further details on the procedure and the scripts used for these analyses are provided in the supplementary info.

For the ‘RMSD/time’ analysis, see Figure 3b) for reference. Here, we plot the RMSD to the closed helix, represented by the black line, where each time point is colored according to the lowest RMSD state. We apply the RMSD/time analysis only to the simulation corresponding to the end state ligand of interest (λ_11_). By tracking the RMSD relative to the closed helix, we can monitor if the protein opens (by high RMSD with green points) or closes (by low RMSD with purple points). Additionally, we gain some insight on the time required to capture the opening or closing of the binding cavity.

**Figure 3:**
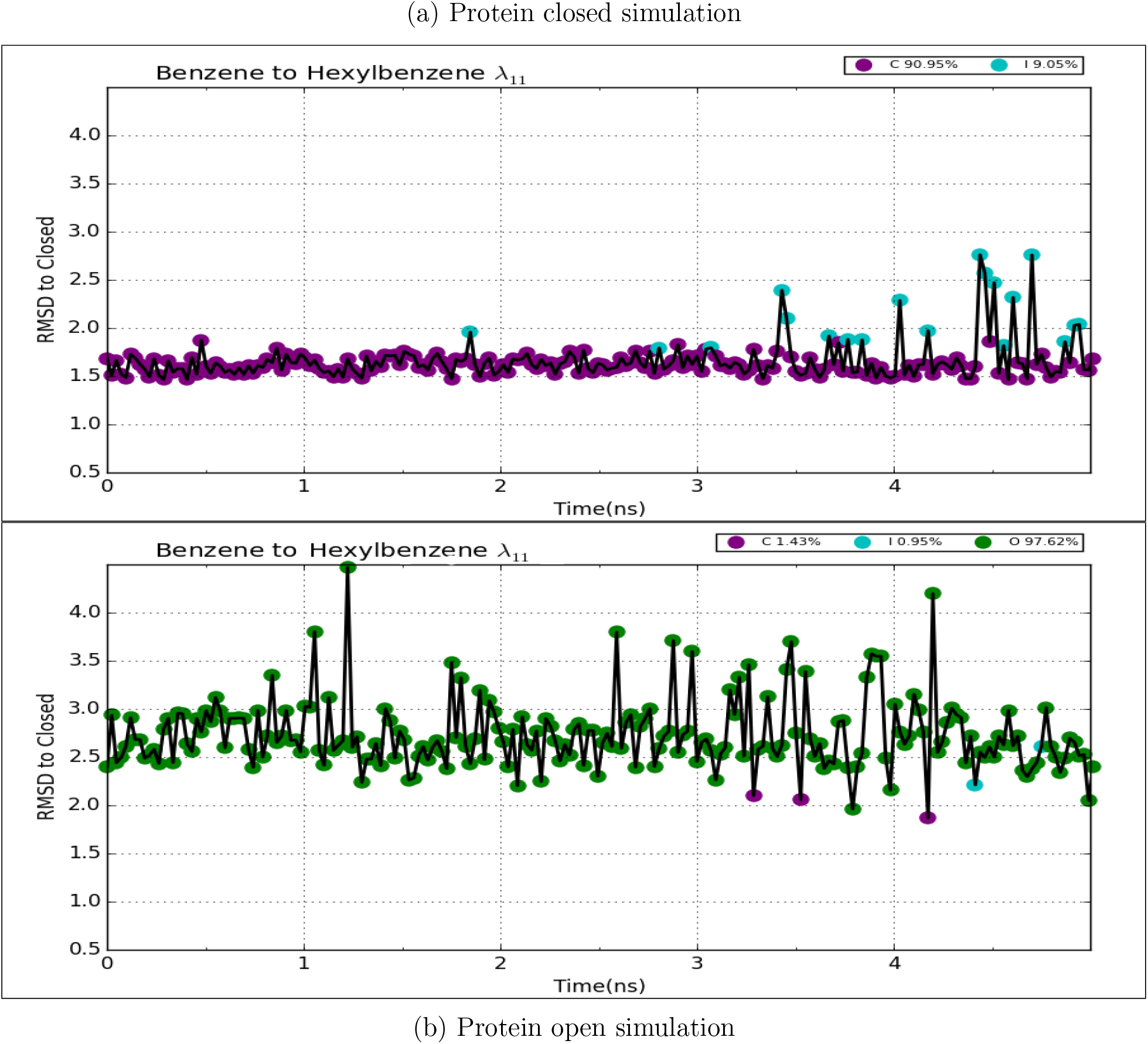
Plotted is the RMSD(Å) relative to the closed helix conformation (black line), where each time point is colored according to the protein state of lowest RMSD to the trajectory. Here, using the default protocol in the transformation of benzene to hexylbenzene, RMSD/time plots correspond to the final end state of hexylbenzene (*λ*_11_). Fig. 3a corresponds to the simulation that began from the protein closed state and Fig. 3b is from the simulation started from the protein open state. The legends indicate the percentage of sampling of each protein conformational state from the trajectory.

It is important to note that by restraining our analysis to only the end-state replica, we limit our ability to completely view the effects of coordinate swapping during replica exchanges. We address this limitation by analyzing all replicas in what we call ‘Color maps’, see Figure 4b for an example. Essentially, our ‘Color map’ analysis is the same as our ‘RMSD/time’ plots but without the RMSD line plot. In other words, we color time points according to the protein state of lowest RMSD and do this for all replicas but do not track the RMSD relative to the closed state. Through a collective view of all replicas, we gain a better perspective of the overall protein conformational sampling and the states they occupy over each replica’s separate trajectories. Using color maps, it becomes visually easy to see if intermediate—higher temperature—lambda windows are able to sample, say the open state, and if this leads to an enhancement in sampling at the end states via replica exchange.

**Figure 4:**
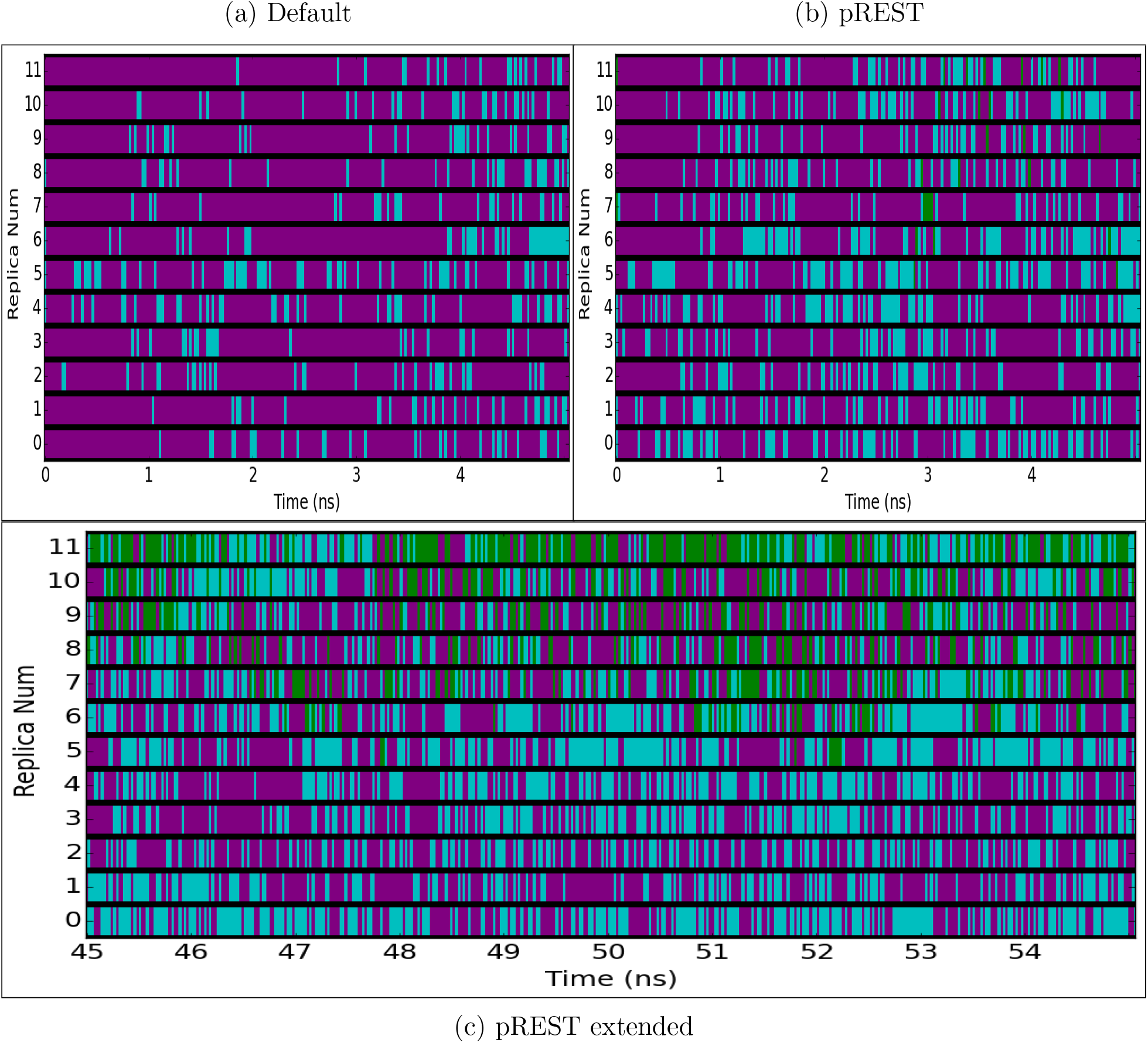
Color maps of simulations starting from the protein closed state for the benzene to hexylbenzene alchemical transformation. Lines at every frame are colored accordingly to the protein state of lowest RMSD (purple-closed, cyan-intermediate, green-open). Fig. 4a corresponds to simulations using the default protocol. Fig. 4b represents simulations using the modified REST region protocol (pREST). Fig. 4c illustrates the enhancement in protein conformational sampling through extending the pREST simulation time up to 55ns.

## 3 Results

### Calculated free energies depend strongly on starting protein conformation

Using the default FEP/REST methodology,^28^ we find calculated free energies significantly depend on the protein starting conformation, especially for large perturbations (i.e. opening the cavity from the closed state). To illustrate this, we begin our molecular dynamics simulations both from the protein closed and open conformations then perform alchemical transformations to ligands that occupy another protein conformational state. For example, in the alchemical transformation of benzene to hexylbenzene—starting from the protein closed state—we expect to see opening of the binding cavity when the ligand is in the fully interacting hexylbenzene state. In this study, we demonstrate using the default FEP protocol settings of a 5ns simulation time and REST region selection does not generate adequate sampling of the motion in the F-helix and does not eliminate the dependence on the initial protein state.

### Closed-Open Ligand Transformations

An examination of the largest alchemical transformation, benzene to hexylbenzene, clearly highlights the sampling challenges faced when using the default FEP/REST protocol. From experimental data of ligand occupancies (Table 1), we expect in our simulations of hexylbenzene to see the protein primarily in the open state over the closed state. Instead, we find the protein remains trapped in its initial conformational state whether we start from closed (Fig 3a) or open (Fig 3b) over the course of the 5ns simulation. From the protein closed simulation, the protein only briefly samples the intermediate state around 3ns but never enters the open conformation. As the protein tries to accommodate hexylbenzene and enter its preferred open state, protein-ligand strain results, yielding a positive value for ΔΔ*G_calc_*(+4.13 kcal/mol). On the other hand, in the protein open simulations, the protein already begins in its preferred state for hexylbenzene and stays only in this open state. As expected, the ΔΔ*G_calc_* is negative(−0.61 kcal/mol) as there is no occurrence of large protein-ligand strain in order to open the cavity. By remaining trapped in the initial state, we under-sample the open state if we begin from the closed state or over-sample it if we begin from the open state. Ultimately, we arrive at two very different relative free energies values, where the inconsistency is as large as +4.74 kcal/mol for the same transformation of benzene to hexylbenzene.

In the overall set, we similarly observe protein closed simulations to yield positive free energies and negative for protein open simulations. In turn, we find the overall inconsistency to be very high with a RMSI of +4 kcal/mol (Fig 5a, Table S4). Clearly, despite the use of implemented default FEP/REST protocol, we are unable to get sufficient sampling in the protein within the standard 5ns time frame. This is perhaps unsurprising since in the default calculation, only the perturbed ligand R-group is added to the enhanced sampling region. Instead, we encounter sampling problems as the protein remains in its initial conformational state throughout the simulation. As a result, our calculated free energies exhibit high dependence on the initial protein configuration which is reflected by the large RMSI.

**Figure 5:**
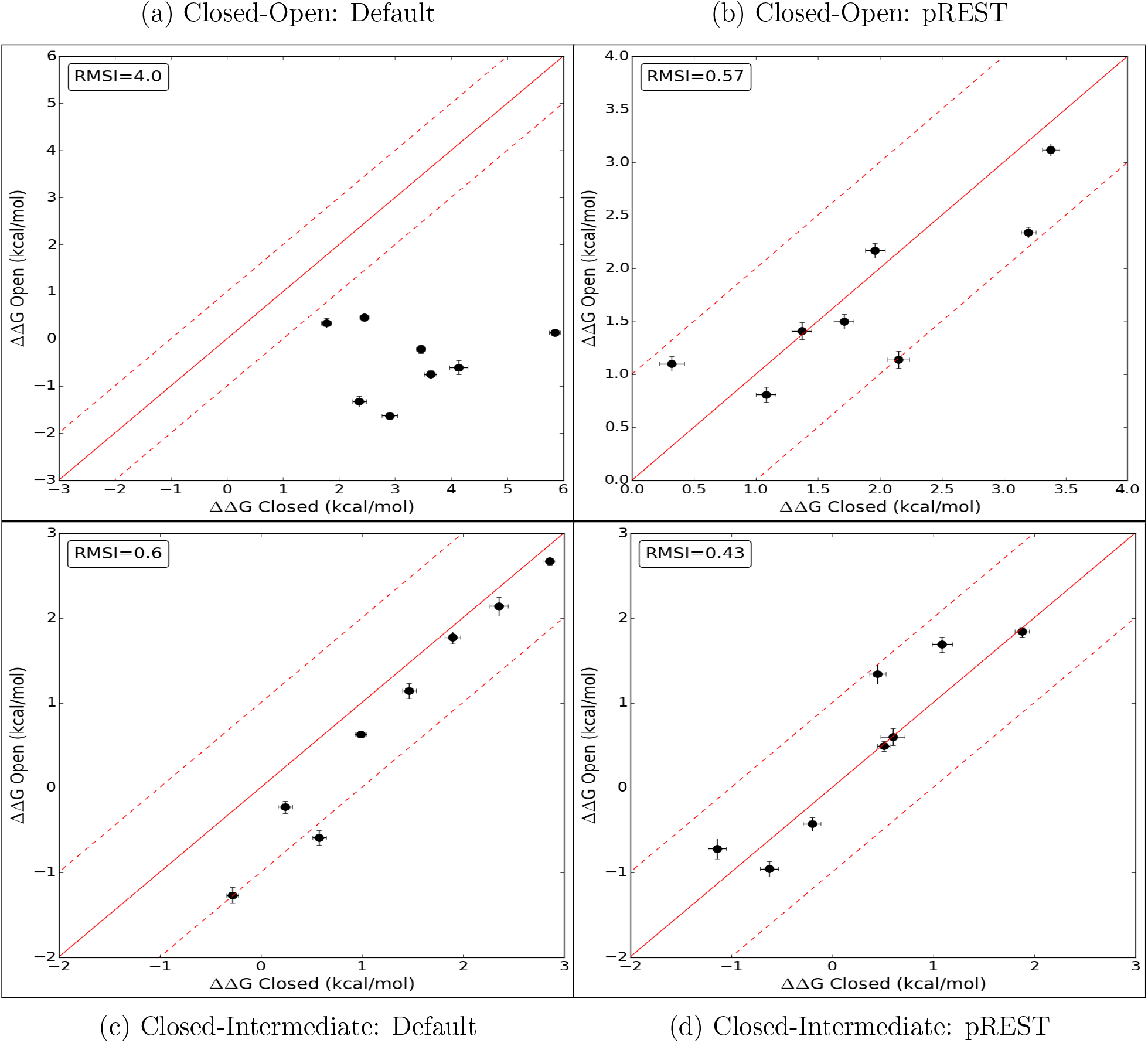
ΔΔ*G_calc_* from MD simulations beginning from the protein closed versus open state. Fig. 5a plots relative free energies obtained using the default protocol from the ‘closed-open’ alchemical transformations set, yielding a RMSI of 4.0 kcal/mol. Fig. 5b are the final computed free energies with simulations carried out to 55ns using pREST, giving RMSI of 0.57 kcal/mol. Fig. 5c plots relative free energies from the ‘closed-intermediate’ set using the default protocol which gives an RMSI of 0.6 kcal/mol. Fig. 5d are free energies with simulations carried out up to 25ns using pREST, giving RMSI of 0.43 kcal/mol. Numerical data for each plot can be found in Tables S1, S3, 4, and SS6.

### Closed-Intermediate Ligand Transformations

In the case of closed-intermediate alchemical transformations, we find that the calculated free energies still have some (albeit much smaller) dependence on the initial protein conformation, using the default protocol. For this set of alchemical transformations, we find the RMSI to be +0.60 kcal/mol (Fig 5c, Table S1). Considering this set involves a smaller protein conformational change and smaller perturbations to the ligand, it is unsurprising to find the RMSI to be much smaller than our closed-open transformation set.

Although, the collective RMSI for closed-intermediate transformations falls in the acceptable range of less than 1 kcal/mol, we can still see a dependence on the initial protein configuration by viewing transformations involving butylbenzene. For these cases in particular, we observe the same pattern of protein closed simulations yielding positive free energies and negative for protein open simulations. We do not see this pattern for transformations with sec-butylbenzene as it does not partially occupy the open state, unlike butylbenzene (Table 1). Through this observation, we demonstrate further that the default protocol does not completely eliminate the free energy dependence on the protein starting conformation, even for smaller perturbations. Although, the level of discrepancy (0.6 kcal/mol) is quite small for this set.

### Experimental Ligand Transformations

Now, when we compare ΔΔ*G_calc_* against ΔΔ*G_exp_*, we find that simulations starting from the protein closed conformation are further from converging to ΔΔ*G_exp_* than when starting from the protein open conformation. Here, we calculate the RMS-’Error’ with experiment and find the RMSE for protein closed simulations to be +1.0 kcal/mol and +0.58 kcal/mol with protein open (Fig 6a, Table S7). Our total RMSI falls within our acceptable range at 0.68 kcal/mol. However, the fact that protein open simulations are much closer to ΔΔ*G_exp_*, once again does demonstrate our calculated free energies depend on the initial protein state. Unsurprisingly, the relatively larger RMSE seen for protein closed simulations primarily comes from transformations involving butylbenzene. Evidently, we find the simulations involving butylbenzene remain trapped in their respective starting conformations, resulting in inadequate sampling in the protein closed simulations (Fig. 7a) versus the protein open simulations (Fig. 7b). Despite performing much smaller alchemical transformations, this shows we still encounter some sampling problems that result in ΔΔ*G_calc_* that depend on the initial protein conformation, evident when comparing to ΔΔ*G_exp_*.

**Figure 6:**
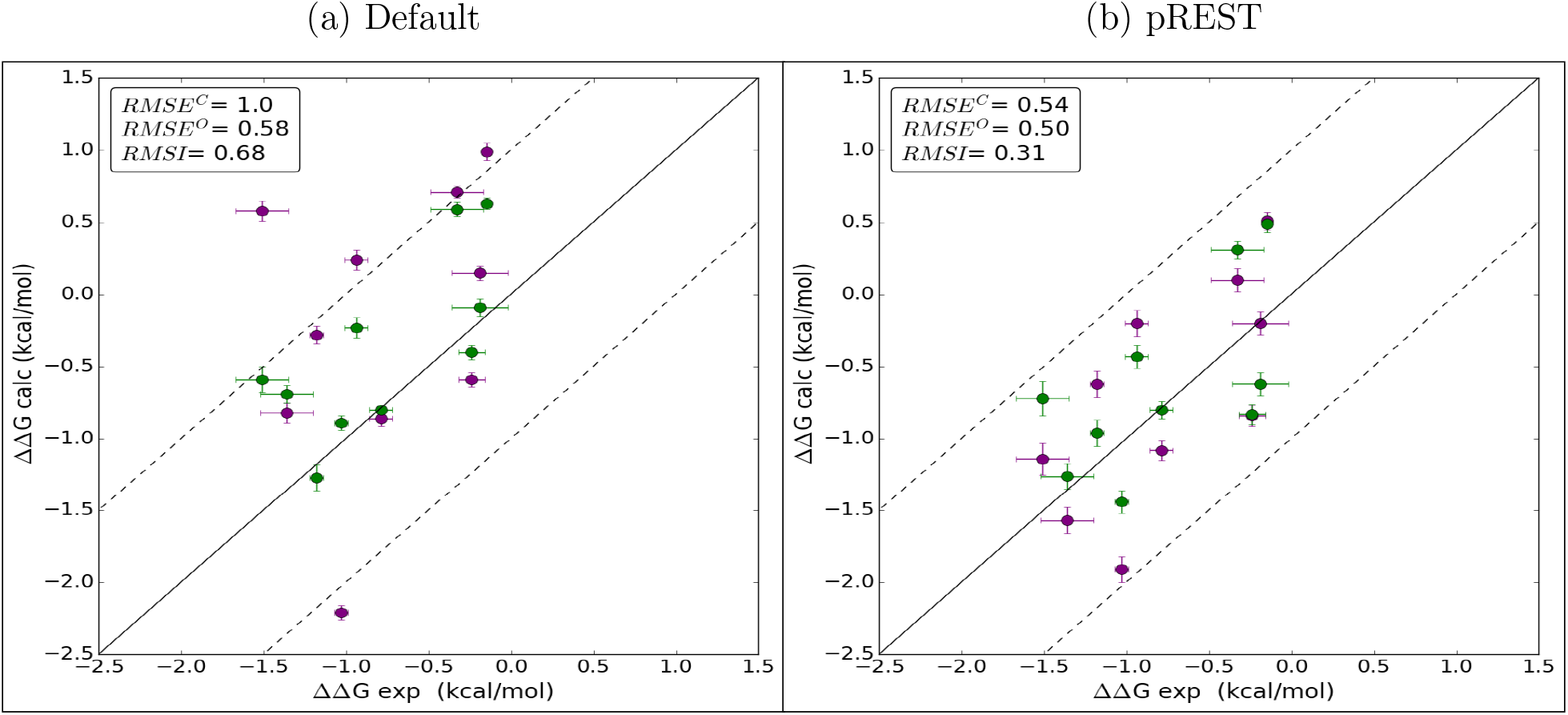
ΔΔ*G_calc_* from MD simulations beginning from the protein closed (purple) and open (green) state compared against ΔΔ*G_exp_*. Fig. 6a plots free energies obtained using the default protocol which gives a 1.0 kcal/mol and 0.58 kcal/mol RMSE for protein closed and open simulations, respectively. Total RMSI using the default protocol is 0.68 kcal/mol. Fig. 6b plots the final free energies after applying pREST, yielding an RMSE of 0.54 for protein closed and 0.50 kcal/mol for protein closed simulations. Total RMSI using the pREST protocol is 0.31 kcal/mol. Numerical data for each plot can be found in Tables S7 and S8.

**Figure 7:**
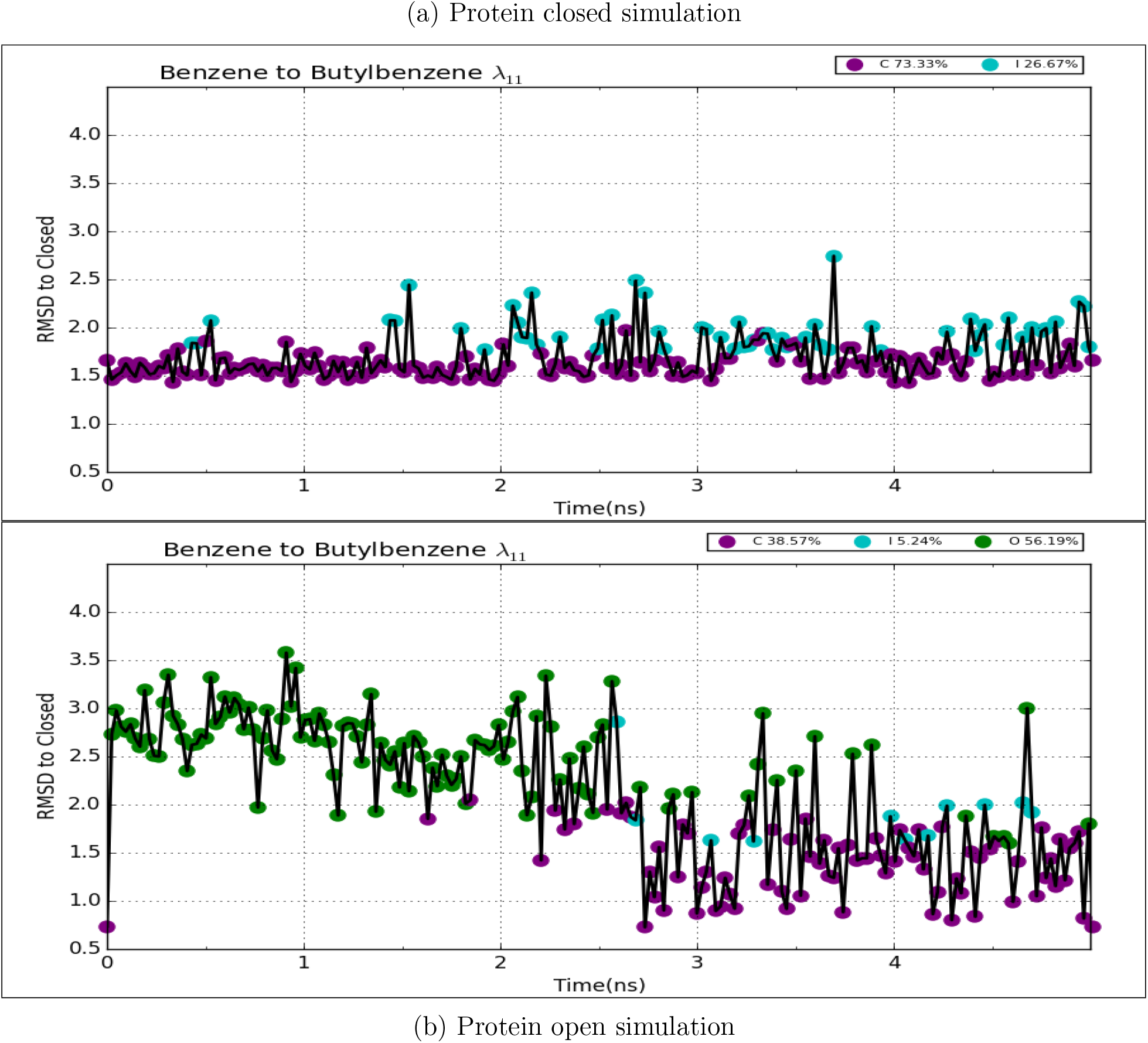
RMSD/time plots correspond to the final end state of butylbenzene (λ_11_) while using the default protocol for transformation of benzene to butylbenzene. Fig. 7a and Fig. 7b correspond to simulations that were started from the protein closed or open state, respectively. The legends indicate the percentage of sampling of each protein conformational state from the trajectory.

### Including protein residues into the REST region (pREST) improves sampling

Primarily, we encounter major sampling problems when we begin our simulations from the protein closed state and attempt a mutation which should result in the binding cavity opening. In order to facilitate protein motion, thereby enhancing protein sampling, we included 3 key residues spanning the F-helix region into the REST region, which we will denote simulations using this with ‘pREST’ (Fig 2a). By expanding the REST region, we are able to drive the F-helix out its initial state trap by locally heating up key regions and thereby reduce our sampling problem.

To demonstrate the REST improvement over the default protocol, we return to the case of benzene to hexylbenzene. Here, we show the facilitation of the helix motion by first referring to Figure 3a which shows that there is no sampling of the open state for the default protocol. Now with pREST, we see a few open state points around 3ns and even a single open point before closing again after our initial step (Fig 8a). Alternatively, we can further illustrate the enhancement of protein sampling by viewing all replicas collectively, using the color maps. In reference to Figure 4a and Figure 4b, we illustrate that there is far less sampling of the intermediate or open protein states in default simulations versus the pREST simulations.

**Figure 8:**
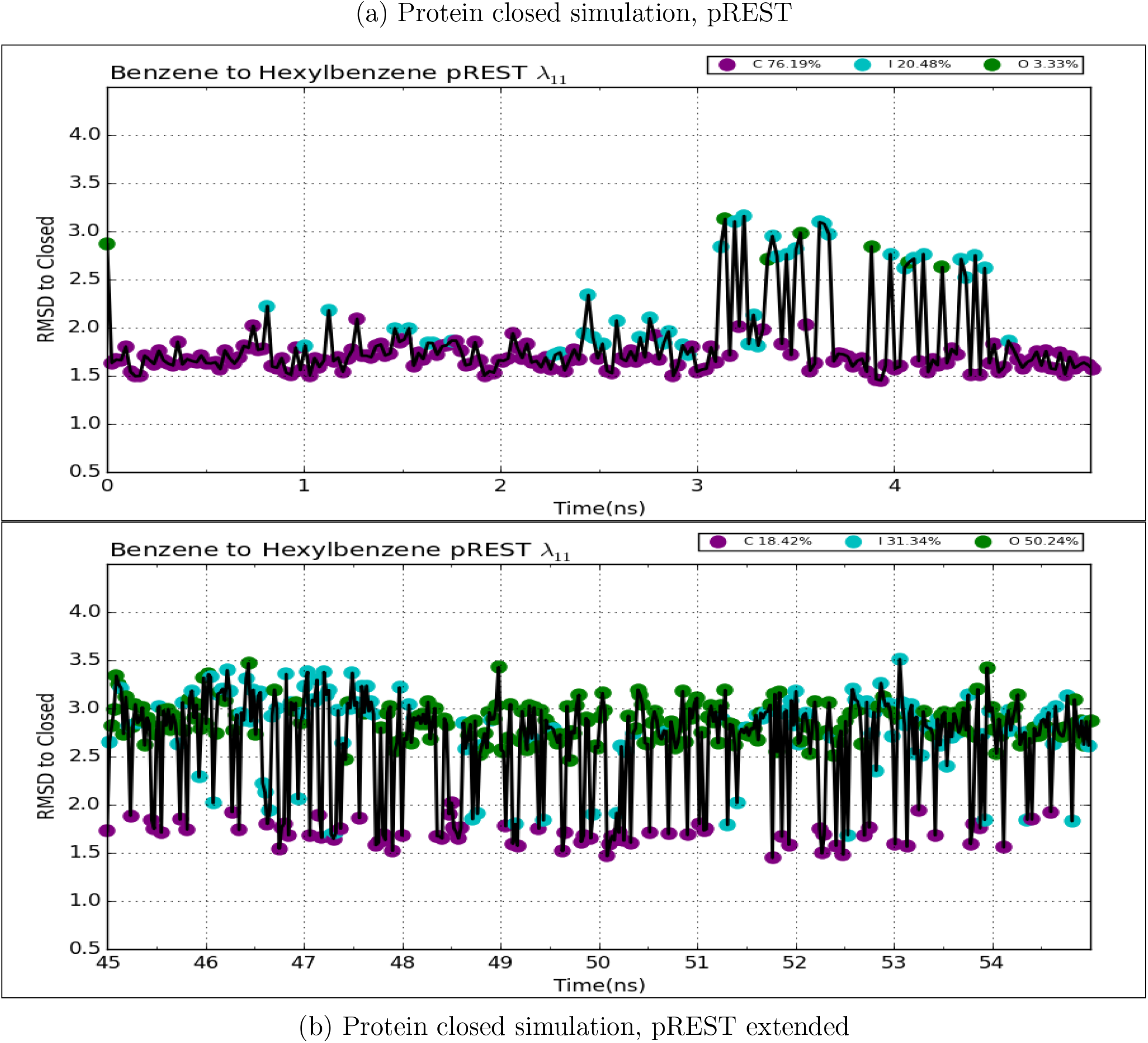
Using the modified REST region (pREST) in the transformation of benzene to hexylbenzene, we plot the RMSD/time corresponding to the hexylbenzene state (*λ*_11_) with the simulation starting from protein closed for the first 5ns (Fig. 8a) and the final 10ns from an extended run up to 55ns (Fig. 8b).

Collectively, we find only some minor improvements in the RMSI for all our closed-open and closed-intermediate transformations while using pREST. For closed-open transformations (Table S5), the RMSI reduces to +2.82 kcal/mol (previously +4 kcal/mol). On the other hand, for closed-intermediate, the RMSI raises slightly to +0.79 kcal/mol (previously +0.60 kcal/mol), which may be due to statistical noise (Table S2). Generally, simulations starting from the closed state had ΔΔ*G_calc_* values that moved towards favorability (i.e. more negative ΔΔ*G_calc_*), while for protein open simulations ΔΔ*G_calc_* values tended towards unfavorability (i.e. more positive ΔΔ*G_calc_*). This is indicative of the fact that pREST is indeed improving sampling, but it is evident that our ΔΔ*G_calc_* are still far from convergence, given the RMSI is still large, especially for closed/open transformations.

### Long simulations enhances protein conformational sampling from more exchanges

Although we see improvements in sampling with pREST, the standard implemented time frame of 5ns clearly is not long enough to gain adequate sampling, particularly if we start from the protein closed state. By running longer, we allow our simulations to perform more exchanges across replicas and thereby allow for better sampling of all conformational states at the relevant end state replicas.

Returning to our most extreme transformation, benzene to hexylbenzene, we have shown pREST alone does not facilitate adequate sampling of the open state (Fig 8a). Now, when we run much longer we see far more sampling of the open protein conformational state in the final 10ns window (Fig 8b). Note that, for closed-open transformations we use the final 15ns to compute the final free energies. We only show the final 10ns in our RMSD/time analysis to avoid overcrowding data points. In viewing all the replicas (Fig 4c), we illustrate the dramatic increase in protein conformational sampling in stark contrast to our previous 5ns simulations (Fig 4b). An additional analysis comparing the protein-ligand contacts between the default and pREST protocol can be found in the Supporting Information (Fig. S1).

By simulating longer with pREST we dramatically increase our sampling of the intermediate and open protein states and almost entirely eliminate the dependence on the initial protein conformational state. For the set of closed-open transformations the RMSI dramatically falls to +0.57 kcal/mol (Fig. 5b, Table S6) and a RMSI of +0.43 kcal/mol for the closed-intermediate (Fig. 5d, Table S3). Similarly, for experimental ligand transformations, our RMSE for protein closed simulations falls to +0.54 kcal/mol and the RMSI reduces to 0.31 kcal/mol (Fig. 6b, Table S8). Now, all our inconsistencies in the final calculated free energies and error from experiment fall within a much more reasonable range of less than +1 kcal/mol.

It is very interesting to note that this overall level of agreement (a typical error of less than 1 kcal/mol) between our calculated values and experiment for the compounds for which we have affinity data – the smaller compounds, for which the inconsistency is relatively low – is actually quite good, better than what was reported in the previous work.^12^ This may be due to the relative simplicity of the ligand modifications (methyl groups only) as well as the high quality of the experimental binding assay, ITC. In contrast, larger apparent errors in other tests may in some cases be due to lower quality experimental data.

## 4 Discussion

In this study, we find that relative free energy calculations for this system can suffer from substantial convergence problems, likely due to the changes being related to the reorganization of a secondary structure element. Although the protein conformational changes in T4 lysozyme (L99A) are extremely localized to a rearrangement of a single helix (Fig 1), we still encounter challenges in sampling, likely due to the slow timescale associated with the reorganization of protein secondary structure. These problems have clear implications for the accuracy of computed relative free energies in these cases. Particularly, we find that calculated relative free energies can depend on the initial protein conformational state by up to 4 kcal/mol.

By looking at alchemical transformations that involve a conformation change in the protein, we show the ΔΔ*Gs_calc_* may be sensitive to the initial protein conformational state when utilizing the default implemented FEP protocol. This sensitivity occurs primarily when the alchemical transformation involves mutating ligands that mainly occupy the closed state (i.e. benzene to propyl) into ligands that occupy the intermediate state (i.e. (sec-)butyl) or, especially, the open state (pentyl/hexyl) (Table 1). By our RMSD analyses, we show the protein remains trapped in its initial state throughout the simulation when using the implemented default protocol. Because it remains trapped, we are unable to adequately sample the necessary protein conformational states and thereby obtain inconsistent ΔΔ*Gs_calc_* depending on whether we start simulations from the open or closed protein configuration. This inconsistency can be large, especially for the larger compounds which bind primarily in the open conformation, for which we do not have experimental binding affinities. However, interestingly, when we start our simulations from the open structure, the lack of adequate sampling of the closed structure does not lead to particularly large errors for the smaller ligands which bind primarily in the closed structure (Table S1).

Without prior knowledge of preferred protein conformational states on ligand binding, we can arrive at very different binding affinity predictions based on the initial protein state being used in the simulation. By starting from the protein closed state and growing the ligand we obtain ΔΔ*G_calc_* values that appear overly positive or unfavorable due to high protein-ligand energy strain and inability to sample the open state, for this system.

However, when we begin with the protein open state our ΔΔ*G_calc_* values appear slightly too negative or favorable, resulting from inability to sample the closed state and not encountering protein-ligand strain. If we only had the crystal structure of the closed protein-ligand complexes, we would blindly conclude that the much larger, open-ligands bind to T4 lysozyme worse than smaller ones. On the other hand, if only the open protein-ligand complexes were available, the opposite would be concluded in that larger ligands are better binders than smaller ligands; experimental data will be needed to determine which is the case.

By including key residues into the REST region and simulating longer, we reduce the ΔΔ*G_calc_* dependence on the initial protein configuration to a more reasonable range of less than 1 kcal/mol. Through expanding the REST region, intermediate lambda windows are able to more easily access the intermediate and open conformations by effectively heating key residues that facilitated protein motion, illustrated in Fig 4b. Further, by simulating longer we allow for more exchanges between replicas, which in turn enhances sampling at our physically relevant end state replica (Fig 4c). With these modifications to the default protocol, we almost completely converge our ΔΔ*G_calc_* to the same value regardless of the starting protein conformation.

Generally, our brute-force approach of simulating longer and multiple trials with varied protein structures is not a desirable or feasible approach, especially in early drug discovery phases. At the industrial level, ligand libraries can be large—driving computational cost exponentially if we simulate longer—or experimental structures can be sparse for new therapeutic protein targets. For future studies, approaches using Markov State Models (MSMs)^64^ can potentially be of great use for identifying discrete protein conformations. MSMs build a representation of the conformational space from batches of short molecular simulations, whereby the discrete states and transition rates between them can be determined in an efficient manner. Utilizing MSMs can thereby provide useful insight on the various protein conformational states before running free energy predictions.

## 5 Conclusions

Overall, we have shown that the presence of kinetically distinct protein conformational states could impact the accuracy of free energy calculations by up to 2-5 kcal/mol, especially so when the R-group modification introduces a severe steric clash with the receptor. It would have been especially challenging as there would essentially be no indicators that the final free energies were sensitive to the initial protein configuration. Only from prior knowledge of the discrete states and by our tedious systematic trials were we able to identify and address the bias in our final calculated free energies. It should be comforting that even without this prior knowledge, good to very good agreement with experiment is obtained for the majority of ligands. Further, one should keep in mind that this study was carefully designed to probe the potential difficulties caused by these conformational changes; most lysozyme ligands studied previously do not induce such significant conformational changes.

Although alchemical free energy calculations have shown tremendous recent successes on a variety of protein targets,^12^ we demonstrate challenges in protein sampling remain. Using T4 lysozyme (L99A) as our simple model system, we highlight sampling problems even from a relatively small (1–3.5Å) and localized single helix rearrangement in response to a series of growing ligands. Through this study, we show using a typical 5ns simulation with only ligand atoms in the REST region, yields free energies that are sensitive to the initial protein conformation. This is especially true for ligand perturbations that induce large protein conformational changes. By longer simulation times and expansion of the REST region to include key protein residues, we were able to get converged predictions and nearly eliminate the depdendence on the starting conformation, even for some challenging cases. This study demonstrates that special attention and care should be exercised when performing alchemical free energy calculations where regions of flexibility surround the binding site. More importantly, prior to performing binding free energy calculations, we present strong evidence on the importance of identifying the occurrence of protein conformational changes upon ligand binding,

## Acknowledgement

N.M.L. thanks Dmitry Lupyan and Joseph Goose for helpful discussions and technical support. Financial support for N.M.L. was provided by Schrödinger and the National Science Foundation Graduate Research Fellowship (DGE-1321846). D.L.M. appreciates financial support from the National Institutes of Health (1R01GM108889-01). L.W. and R.A. are supported by Schrödinger.

---

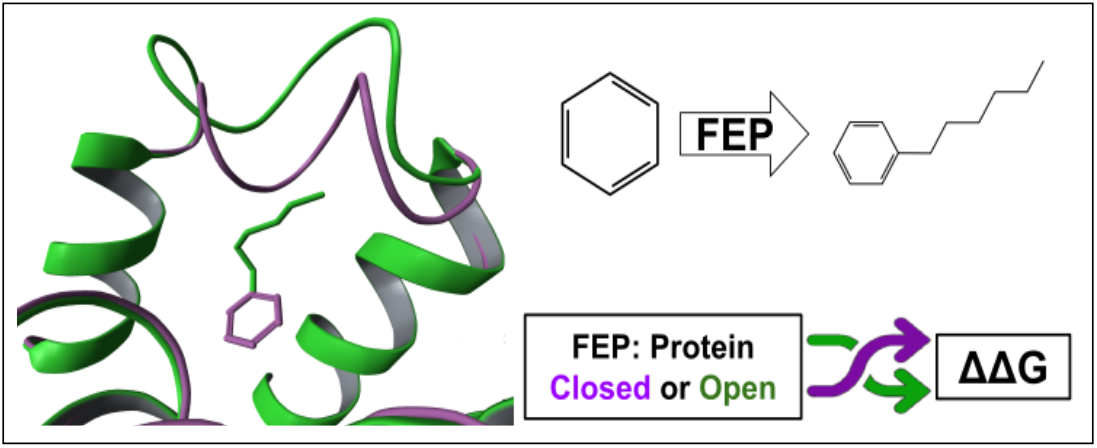
Graphical TOC Entry

